# SALRR: Scalable Analysis of Long-Read RNA-Seq Enables Comprehensive Transcriptome Profiling in Human Brain

**DOI:** 10.64898/2026.08.27.747499

**Authors:** Cedric Kouam, Jackson Mingle, Pilar Álvarez Jerez, Allison Evans, Abraham Moller, Breeana Baker, Cory Weller, Kimberly Paquette, Janet Brooks, Spencer M Grant, Alexis Ayuketah, Melissa Meredith, Joanna Palade, Laksh Malik, Kristen Hise, Jessica Anderson, Robert Harbert, Yilei Fu, Xinchang Zheng, Sonia García-Ruiz, Emil K Gustavsson, Cornelis Blauwendraat, Mina Ryten, Fritz J Sedlazeck, Luigi Ferrucci, J. Raphael Gibbs, Jinhui Ding, Xylena Reed, Mike Nalls, Mark R Cookson, Kendall Van Keuren-Jensen, Elizabeth Hutchins, Miten Jain, Kimberley J Billingsley

## Abstract

Isoform-resolved transcriptomics is fundamental to decoding the molecular complexity of the human brain, yet population-scale long-read RNA sequencing has remained inaccessible due to labor-intensive library preparation, sensitivity to RNA degradation in postmortem tissue, and the absence of integrated, reproducible analysis pipelines. Here we present SALRR (Scalable Analysis of Long-Read RNA-seq), an integrated wet-lab and computational platform designed to overcome these barriers. Automated ONT long-read cDNA library preparation on the Hamilton Microlab NGS STAR platform reduces hands-on time by 67% and enables 24 libraries per operator per day while maintaining performance across RNA integrity values. A modular, Snakemake-based pipeline performs end-to-end processing from ONT signal data to isoform-level quantification, incorporating SIRV spike-in calibration, multi-stage quality control, and stringent isoform validation. Applied to 10 postmortem frontal cortex samples from the North American Brain Expression Consortium, SALRR identified 31,607 high-confidence isoforms from 10,075 genes, including 8,532 novel splice variants absent from GENCODE v49, and complex splicing events systematically missed by short-read sequencing at neurodegeneration-relevant loci, including *GBA1*, *CCNF*, *CHCHD10*, and *TREM2*. All protocols and code are openly available, providing a scalable, community-ready framework for isoform-resolved transcriptomics in neurodegeneration, aging, and complex brain disease.

---

The human brain expresses a greater diversity of transcripts than any other tissue, with alternative splicing, gene fusions, and isoforms arising from repeat-rich regions generating thousands of functionally distinct products ^1^. This transcriptomic complexity is central to neuronal identity, synaptic function, and disease susceptibility ^2,3^, yet it remains poorly characterized at the isoform level: current approaches to quantification obscure transcript-specific regulation and produce systematic errors when isoform switching alters function without changing total gene expression ^4^.

Short-read RNA sequencing has enabled transcriptome-wide profiling at scale, but with read lengths of approximately 200 bp (far shorter than the median human transcript), it cannot resolve full-length isoform structures, establish exon connectivity across long transcripts, or detect complex splice junctions. Long-read RNA sequencing (LR-RNAseq) overcomes these limitations by capturing full-length transcripts in a single read, enabling accurate isoform quantification, novel transcript discovery, and detection of gene fusions and splicing events in repeat-rich regions ^5,6^. Both Oxford Nanopore Technologies (ONT) and Pacific Biosciences (PacBio) support full-length cDNA sequencing; ONT additionally offers direct native RNA sequencing for epitranscriptomic analysis and accommodates a broader range of molecule lengths, while PacBio delivers higher per-read accuracy ^7^. Comprehensive benchmarking has established long-read approaches as substantially superior to short-read methods for isoform-level analyses ^8^.

LR-RNAseq has demonstrated the ability to link noncoding splicing variants to altered protein isoforms ^9,10^ and, in combination with multi-omic data, to identify disease mechanisms and therapeutic targets inaccessible to gene-level approaches ^11,12^. In postmortem brain tissue specifically, long-read methods have uncovered extensive novel isoform diversity relevant to aging and neurodegeneration^13,14^, underscoring the scale of biologically important information that short-read studies miss. Yet population-scale implementation in human postmortem tissue remains severely limited by three barriers that have not been simultaneously addressed: labor-intensive and low-throughput library preparation protocols; RNA degradation in postmortem samples; and the absence of standardised, end-to-end analysis pipelines. Current tools for isoform detection and quantification vary widely in precision, sensitivity, and computational efficiency, and typically require integration of multiple software packages to achieve comprehensive results^15^.

Realizing the potential of LR-RNAseq for population-scale brain studies requires workflows that are simultaneously scalable, cost-effective, robust to variable RNA quality, and paired with reproducible and integrated analysis pipelines. We have developed SALRR (Scalable Analysis of Long-Read RNA-seq), a platform that meets these requirements. SALRR combines an automated, high-throughput ONT cDNA library preparation workflow on the Hamilton Microlab NGS STAR platform (optimized for postmortem human brain tissue and tolerant of variable RNA integrity) with a modular, Snakemake-based computational pipeline that encapsulates basecalling, read re-orientation and trimming, genome mapping, transcriptome reconstruction, and gene and isoform-level quantification in a single reproducible workflow. Together, these advances enable isoform-resolved transcriptomics for the human brain at a scale and robustness not previously achievable.

## RESULTS

### An integrated platform for scalable long-read RNA sequencing from postmortem human brain

We developed SALRR, an integrated wet laboratory and computational platform for population-scale transcript isoform analysis in postmortem human brain tissue (Fig. 1). The laboratory component comprises optimized RNA extraction protocols for cultured cells and frozen frontal cortex tissue, followed by an automated ONT cDNA library preparation workflow on the Hamilton Microlab NGS STAR liquid handling platform. This is paired with a modular, Snakemake-based computational pipeline that processes ONT POD5 data through basecalling, full-length read identification, reference alignment, transcript reconstruction, and isoform-level quantification (Supplementary Fig. 1). We applied this platform to 3 cultured iPSC samples and 10 frontal cortex samples from neurologically normal individuals enrolled in NABEC. This dataset provides substantially deeper transcript-level resolution than has previously been available for postmortem brain, capturing full-length isoform structure and novel transcript discovery at a granularity inaccessible to short-read approaches. All data and processing pipelines are publicly available.

**Fig. 1:**
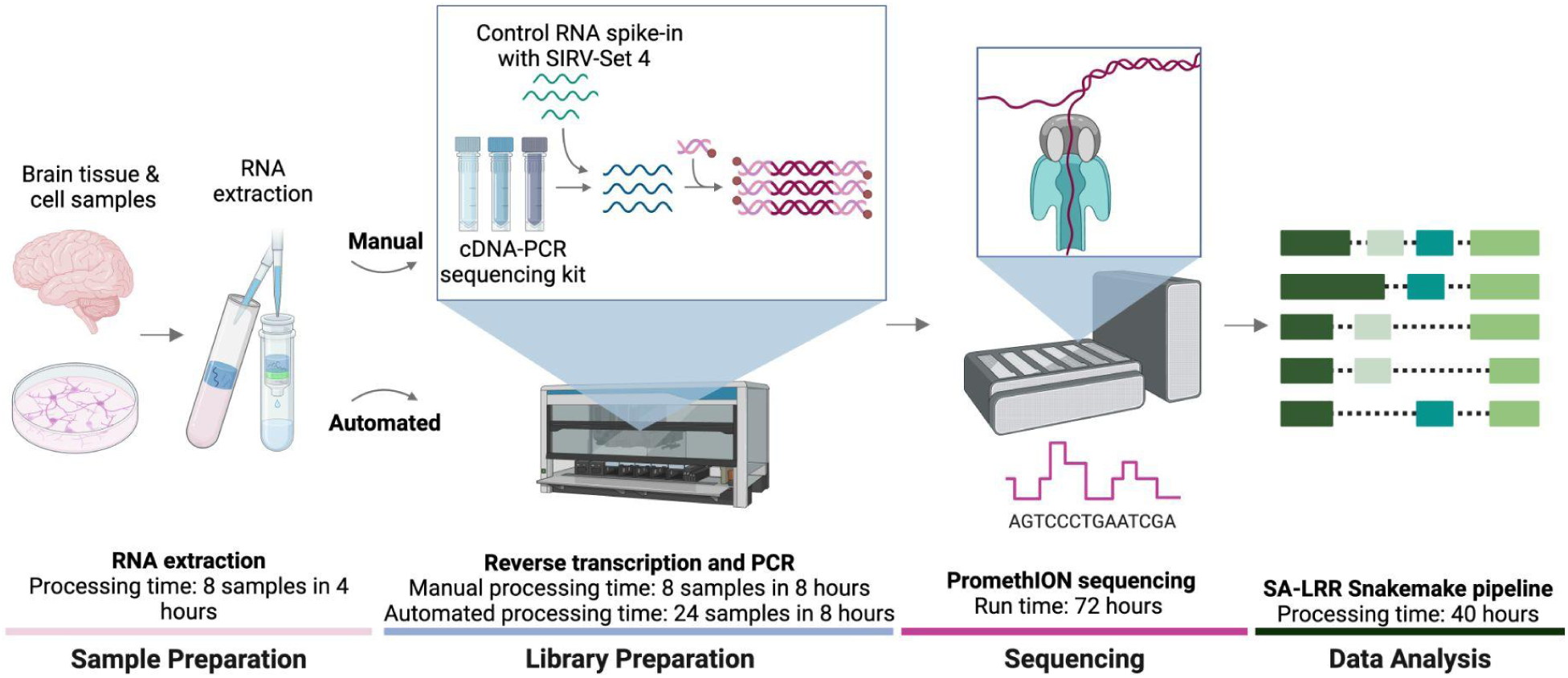
SALRR enables scalable, automated long-read RNA sequencing from postmortem human brains. Schematic of the SALRR wet-lab workflow, showing manual and automated library preparation routes, SIRV spike-in addition, PromethION sequencing, and computational processing. Processing times per stage are indicated. Data analysis processing time (∼40 hours) reflects runs executed on the NIH Biowulf HPC cluster (detailed computational resources and sample runtimes provided in Supplementary Table 6). Created in BioRender. Billingsley, K. (2026) https://BioRender.com/59yhkkt

**Supplementary Fig. 1:**
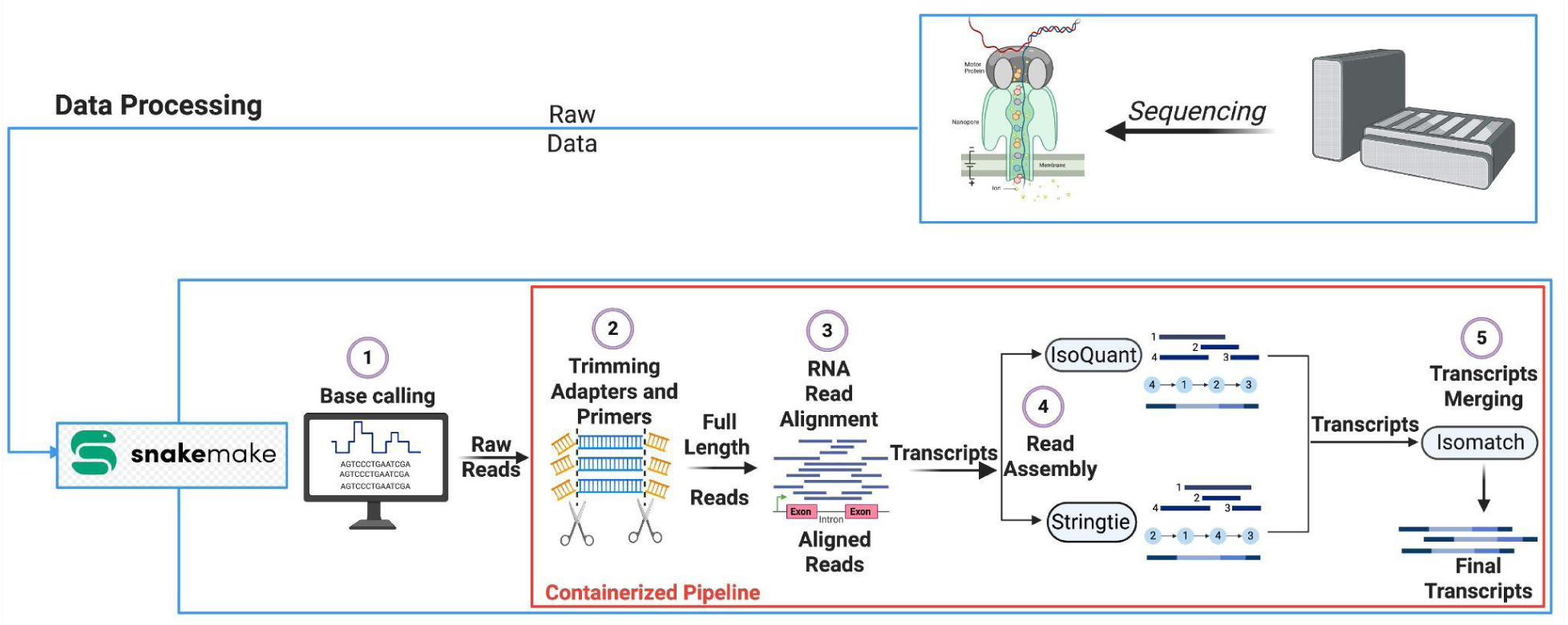
Architecture of the SALRR computational pipeline. Modular Snakemake-based pipeline for end-to-end processing of Oxford Nanopore long-read RNA sequencing data from raw POD5 signal to isoform-level quantification. Input POD5 files are basecalled using Dorado v1.2.0 with the super-accuracy model, with poly(A) tail estimation performed concurrently. Unmapped BAM files are converted to FASTQ format using SAMtools v1.18 and processed through Pychopper v2.7.10 for full-length cDNA read identification, rescue, and quality filtering (minimum Q10, minimum 300 bp). Read-level quality metrics are collected at each stage using Cramino. An optional contamination screening step using FastQ Screen with minimap2 alignment can be enabled or disabled within the pipeline configuration. When SIRV spike-in controls are present, trimmed reads are first aligned to the SIRV reference genome using minimap2 v2.28; unmapped reads are then realigned to the human reference genome GRCh38. When SIRV controls are absent, reads are aligned directly to GRCh38. Alignment quality control is performed using SAMtools, Cramino, and mosdepth v0.3.3. Only primary alignments with a mapping quality score greater than 40 are retained. Transcript reconstruction is performed in parallel using IsoQuant v3.10.0 and StringTie v2.2.1, and resulting isoform catalogues are merged using Isomatch v0.4.0 to generate a consensus set of transcripts. Novel isoforms are classified by structural category using SQANTI-like classification relative to GENCODE release 49. All modules, branching logic, intermediate file formats, and QC checkpoints are shown. Created in BioRender. Billingsley, K. (2026) https://BioRender.com/59yhkkt

### PromethION sequencing delivers consistent, high-quality output

Across 3 iPSC and 10 NABEC brain tissue samples sequenced on the PromethION, runs yielded a mean of 68.8 million reads (170.4 Gb; Supplementary Table 2), a median read length of 2,064 bp, a median quality score of Q24.91 (Supplementary Fig. 2), and consistently high mapping efficiency to GRCh38 (mean 99.93% uniquely mapped reads; Fig. 2a).

**Fig. 2:**
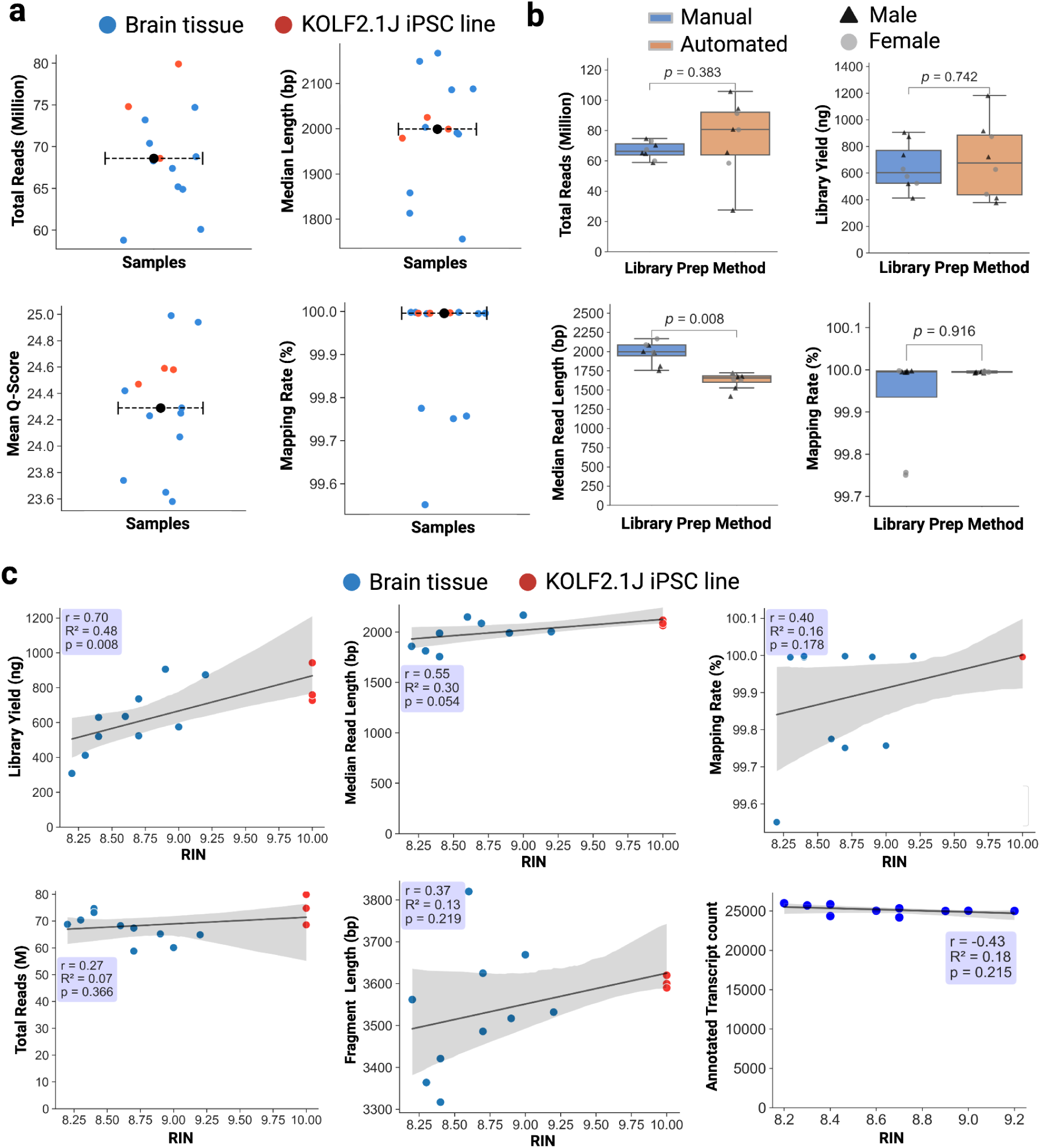
Benchmarking manual and automated library preparation performance across sample types and RNA quality. **a,** Per-sample sequencing metrics. Total reads (millions), median aligned read length (kb), mean Q score, and GRCh38 mapping rate are shown for all samples. Points indicate individual samples; black points and dashed lines indicate the median. **b,** Library preparation metrics for manual versus automated workflows using matched RNA samples. Library yield, median aligned read length, total reads (millions), and mapping rate are shown. Each point represents an independent preparation; boxes show median and interquartile range; p-values from two-sided Wilcoxon signed-rank tests. **c,** SALRR performance across a range of RNA integrity numbers (RIN). Library yield, total reads, median aligned read length, mapping rate, fragment length, and annotated transcript recovery are shown as a function of RIN. Lines indicate linear regression fits; shading indicates 95% confidence intervals. n = 13 libraries (10 brain, 3 KOLF2.1J iPSC); annotated transcript recovery is shown for brain samples only (n = 10).

**Supplementary Fig. 2:**
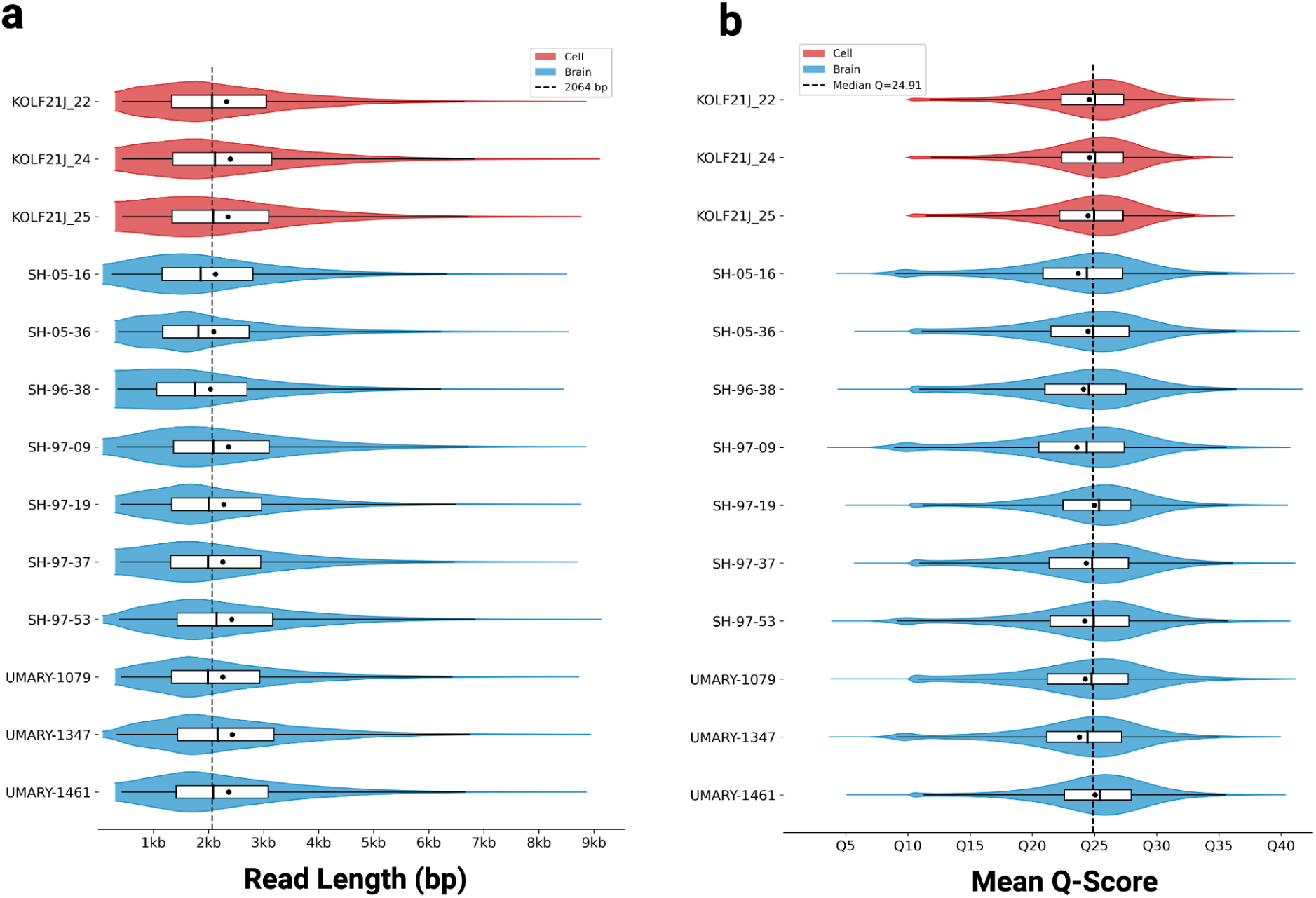
Sequencing quality control metrics for mapped reads. Violin plots with overlaid boxplots show per-sample distributions of a, read length (bp) and b, mean Q-score. Samples are coloured by type: red, KOLF2.1J iPSC; blue, brain tissue. Dashed lines mark the overall median read length (2,064 bp) and the overall median Q-score (24.91).

### Automation increases throughput, reproducibility, and scalability

A key barrier to population-scale LR-RNAseq has been the labor intensity and low throughput of manual library preparation protocols. We therefore automated cDNA library preparation end to end, from RNA input to sequencing-ready library in a single unattended run, and compared it against the manual protocol using matched frontal cortex RNA (Fig. 2b). Automation reduced hands-on time by 67% and increased throughput from 8 to 24 libraries per person per day, with no significant difference in library yield (p = 0.742) or mapping rate (99.99%, p = 0.916) and only a modest, though statistically significant, reduction in median aligned read length (2.0 kb manual vs. 1.7 kb automated, p = 0.008); both represent a ∼5-fold increase over previous benchmarks^13^ (345 bp). The automated protocol therefore increases throughput and scalability while maintaining data quality. RNA degradation is a pervasive challenge in postmortem brain studies, where RNA integrity numbers (RINs) frequently fall below thresholds recommended for standard protocols. Across libraries spanning RIN 8.2–10 (Fig. 2c), total library yield correlated positively with RIN (Pearson r = 0.70, p = 0.008), while read throughput (mean 68.85 million reads) and annotated transcript counts remained stable across the RIN range (r = 0.27, p = 0.366; and, for brain samples only, r = -0.43, p = 0.215), indicating that transcript reconstruction is robust to the RNA quality typical of postmortem tissue.

### SALRR enables accurate gene and transcript level quantification

To validate SALRR, we benchmarked 10 NABEC brain samples against matching short-read data from the same cohort, restricting analysis to shared reference genes and transcripts (Fig. 3a,b). SALRR TPM estimates correlated strongly with short-read data at both the gene level (Pearson r = 0.742, Spearman ⍴ = 0.721; Fig. 3a) and the transcript level (r = 0.76; Fig. 3b), compared with previous benchmarks (r² = 0.75 and 0.57; Glinos et al^17^). Long-read TPM values are systematically higher because fewer genes are detected per sample (mean 21,935 vs. 32,917), distributing the fixed 10⁶ TPM sum across fewer targets; this compositional shift preserves relative gene rankings, confirmed by the strong rank correlation. SALRR expression estimates for ERCC controls within the SIRV-Set 4 spike-in mix correlated strongly with known input concentrations across brain (r = 0.939) and cell (r = 0.943) libraries (Fig. 3c, Supplementary Fig. 3a,b, Supplementary Table 5), confirming accurate quantification of highly expressed genes and transcripts.

**Fig. 3:**
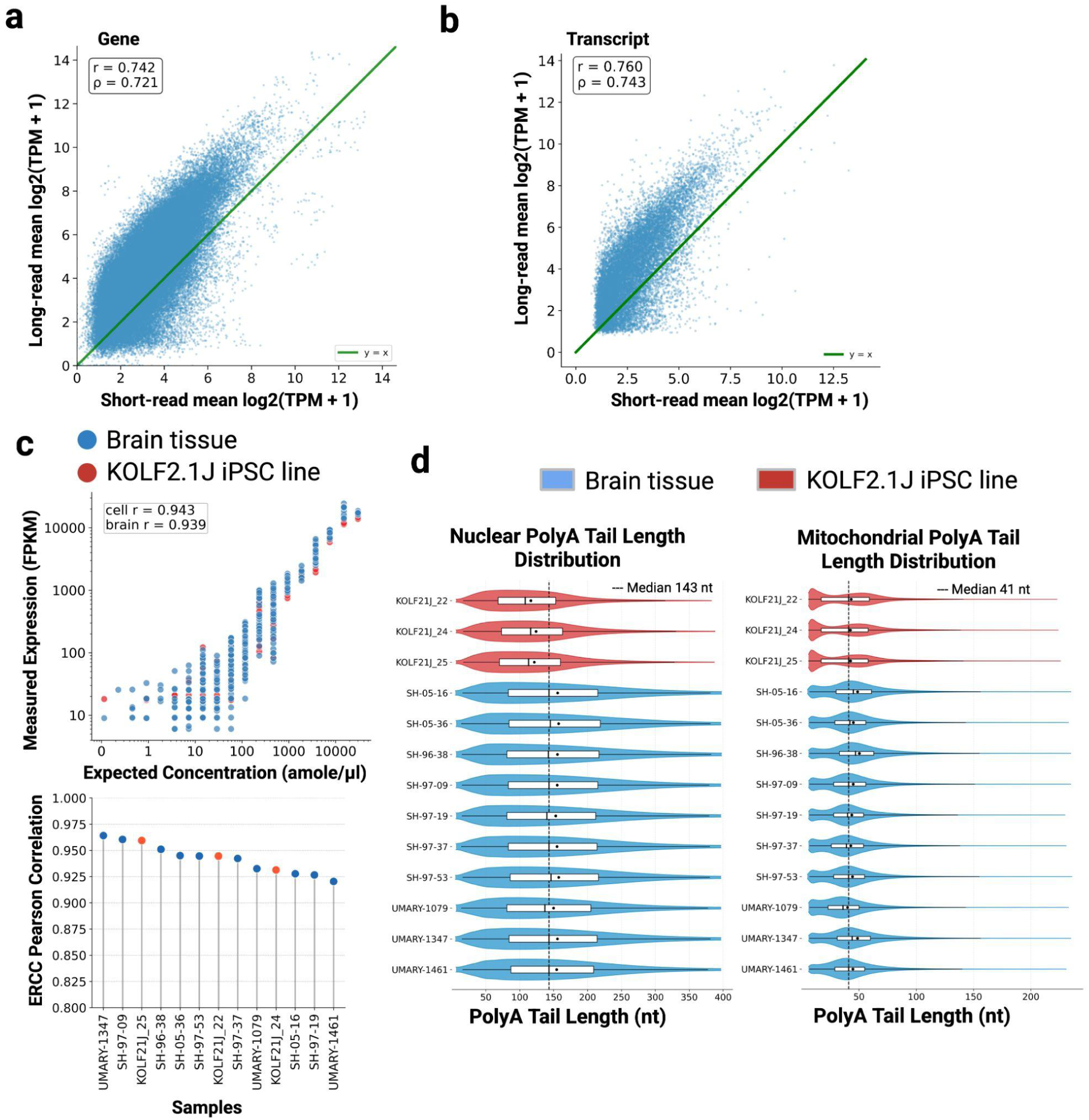
SALRR quantification agrees with short-read RNA-seq and resolves transcript-level detail. **a,** Correlation between gene-level expression estimates from SALRR long-read data and matched short-read RNA-seq data from the same NABEC frontal cortex samples. Each point represents one expressed reference gene quantified by both methods; solid green line indicates y = x identity (Pearson r = 0.742, Spearman ⍴ = 0.721). **b,** Correlation between transcript-level expression estimates from SALRR and matched short-read RNA-seq data for the same samples (Pearson r = 0.760, Spearman ⍴ = 0.743). **c,** ERCC spike-in recovery across iPSC (n = 3) and brain tissue (n = 10) samples. Top: Observed expression versus expected concentrations (r = 0.943 for cell lines, r = 0.939 for brain tissue). Bottom: Per-sample Pearson correlation values for ERCC spike-ins. **d,** Nuclear and mitochondrial poly(A) tail length distribution across all 10 brain samples and 3 cell line samples estimated during basecalling with Dorado. Violin plots show per-sample distributions, with median poly(A) tail lengths of 143 nt (nuclear) and 41 nt (mitochondrial) marked by dashed lines, consistent with published human brain polyadenylated RNA estimates^16^. Density near the lower threshold reflects the detection boundary of the Dorado basecaller (∼6 nt) rather than true short tails.

Long-read sequencing further provides transcript features absent from conventional short-read data. Poly(A) tail length, estimated during basecalling, showed a median of 143 nucleotides for nuclear transcripts, matching published mammalian estimates^16^ while mitochondrial transcripts were markedly shorter (41 nt)^18^, consistent with the known short poly(A) tails of human mitochondrial mRNAs (Fig. 3d). Read-level assignment metrics were also consistent with comparable human brain long-read cDNA datasets: a mean of 45% of reads were uniquely assigned to individual isoforms and 15% spanned the full length of their assigned transcript (Supplementary Fig. 3c), in line with the 42% and 17% reported by Aguzzoli Heberle et al^13^, with low sample-to-sample variation indicating consistent library quality. Gene body coverage analysis supported transcript completeness, showing more uniform coverage, better recovery at the 5’ end and greater 3’ retention in long-read versus short-read data (Supplementary Fig. 3d).

**Supplementary Fig. 3:**
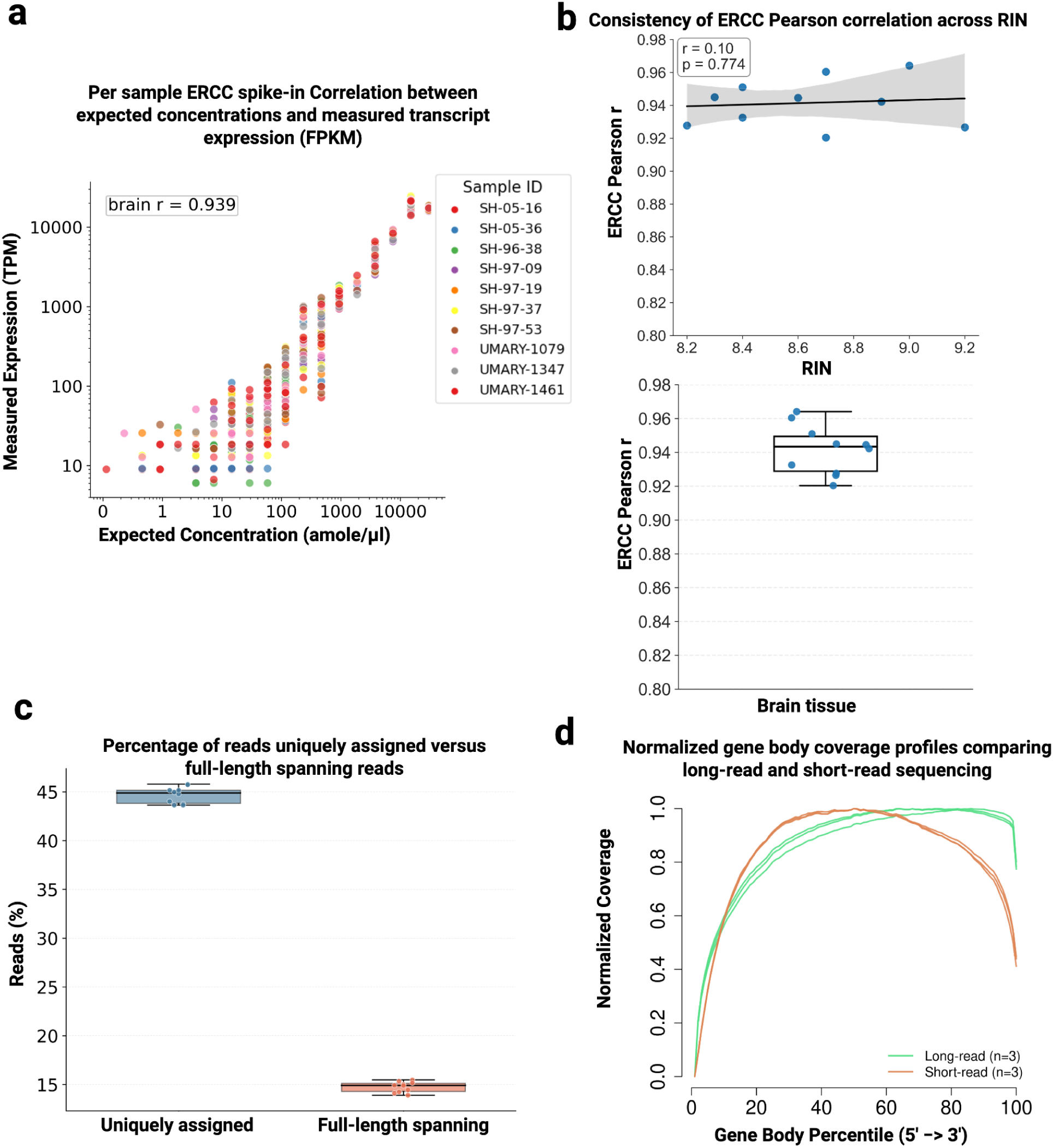
Assessment of quantitative accuracy, coverage uniformity, and alignment efficiency across samples. **a,** Correlation between expected (amole/μl) and observed expression (TPM) of ERCC spike-in controls transcript abundances across all 10 brain samples (r = 0.939). Points are coloured by sample ID. **b,** Stability of ERCC quantitative accuracy across RNA integrity scores. Top: Pearson correlation (r) between expected and observed ERCC abundances plotted against sample RIN (r = 0.10, p = 0.774), demonstrating that quantification accuracy is maintained across variable RNA quality. Bottom: Distribution of per-sample ERCC Pearson correlation values across brain samples (mean r = 0.94, range: 0.92–0.96; points indicate individual samples, box shows median and IQR). **c,** Distribution of sequencing reads categorized by assignment category, comparing uniquely assigned reads (∼45%) against full-length transcript-spanning reads (∼15%). Each point represents an individual sequencing library. **d,** Normalized gene body coverage profiles comparing long-read (n = 3, green) and short-read (n = 3, orange) sequencing across normalized transcript length (5’ to 3’ percentile).

### Comprehensive isoform detection and novel transcript discovery in the human brain

Next, we characterized the transcriptomic landscape of the human postmortem frontal cortex at isoform-level resolution (Fig. 4a). On average, the pipeline detected 36,318 transcripts per sample. Assembly models from IsoQuant and StringTie were filtered to remove low-abundance artifacts (CPM > 1) and merged per sample using Isomatch and GENCODE v49. Because merge models retain transcript structure without cohort-level requantification, counts reflect per-sample detection rather than cohort-wide expression. An average of 64% (23,243) matched GENCODE v49 reference transcripts (full-splice match (FSM) or incomplete-splice match (ISM)) and 36% (13,074) were classified as novel or unannotated, a fraction consistent across samples and dominated by novel-in-catalog (NIC) and novel-not-in-catalog (NNC) transcripts (Fig. 4a,b).

**Fig 4:**
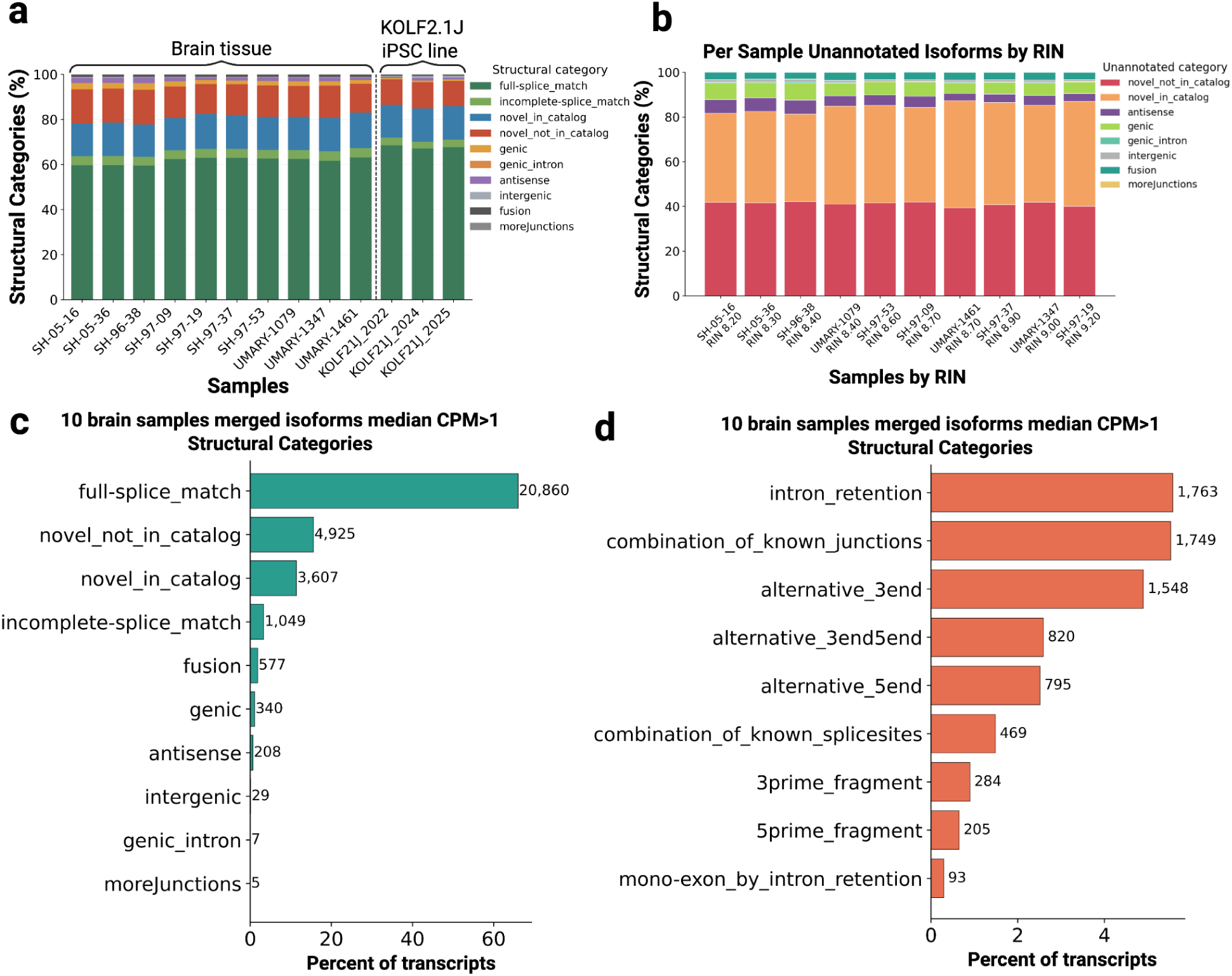
Isomatch structural annotation profiling and novel isoform subcategory distribution of pass-threshold transcripts. **a,** Relative distribution of transcript structural categories classified by Isomatch (using SQANTI-style nomenclature) across individual postmortem brain tissue (n = 10) and KOLF2.1J iPSC line (n = 3) samples. **b,** Relative proportion of unannotated isoform categories across brain samples ordered progressively by RNA integrity number (RIN 8.20 to 9.20), demonstrating consistent structural category representation across variable RNA quality. **c,** Aggregate structural category distribution and absolute transcript counts for high-confidence merged brain isoforms (median CPM > 1 across n = 10 samples). Numbers adjacent to bars indicate absolute transcript counts; full-splice match (FSM) isoforms represent the largest category (20,860 transcripts). **d,** Subcategory breakdown and absolute counts for novel structural classification events (e.g., *intron retention*, *combination of known junctions*, *alternative 3’ end*) among high-confidence filtered brain transcripts (median CPM > 1).

To identify robustly expressed isoforms, we collapsed per-sample models into a unified transcript set using Isomatch, requantified reads against this reference, and applied expression filtering (median CPM > 1 across the 10 samples) adapted from Aguzzoli Heberle et al.^13^; given the high sequencing depth (minimum 40 million mapped reads per sample vs. ∼24 million previously), this threshold provides a stringent filter for high-confidence calls. In total, we identified 31,607 expressed isoforms from 10,075 genes: FSM/ISM isoforms comprised ∼69% (21,909), while novel splice variants (NIC/NNC) accounted for ∼27% (8,532; Fig. 4c), with intron retention, novel combinations of known junctions, and alternative 3’ ends the most frequent novel subcategories (Fig. 4d, Supplementary Table 4). By biotype, protein-coding genes accounted for 85.3% of expressed isoforms, followed by intron retention (5.9%), lncRNA (4.6%), and other biotypes (4.2%) (Fig. 5b); this exceeds the ∼70% reported by Aguzzoli Heberle et al. 13 and 65–82% by Page et al. ^19^ because parent-gene-level assignment groups non-coding transcripts of protein-coding genes (e.g., NMD substrates, processed transcripts) under the protein-coding umbrella. Within this group, 73.6% (19,847) were known FSM/ISM transcripts and 26.4% (7,117) were novel; lncRNA showed a similar split (85.0% known), while intron-retention isoforms were predominantly novel (74.8%).

**Fig 5:**
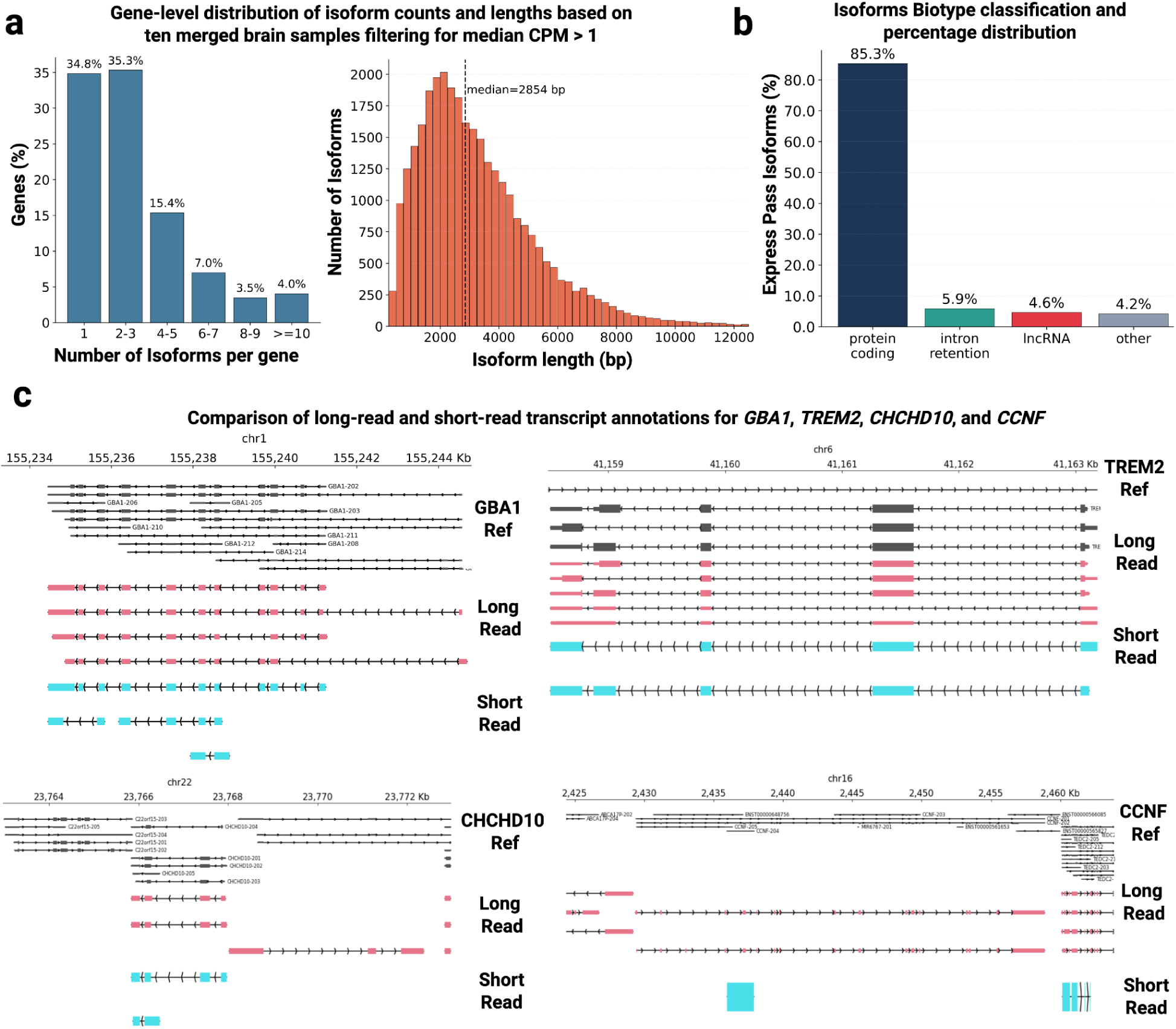
Isoform complexity, biotype classification, and structural resolution at neurodegenerative disease loci. **a,** Left: Distribution of the number of expressed transcript isoforms identified per gene across postmortem brain samples (filtered for median CPM>1 across n = 10 samples). A total of 65.2% of expressed genes produce multiple transcript isoforms (⩾ 2). Right: Isoform length profile (bp) for high-confidence expressed transcripts, with a median transcript length of 2,854 bp (dashed line). **b,** Biotype classification of high-confidence expressed isoforms, dominated by protein-coding transcripts (85.3%), followed by intron retention (5.9%), lncRNAs (4.6%), and other biotypes (4.2%). **c,** Genomic alignment tracks comparing reference transcript annotations (GENCODE, gray/black), long-read SALRR assemblies (pink), and short-read RNA-seq reconstructions (cyan, StringTie v3.0.1) across four neurodegenerative disease-associated loci: GBA1 (chr1), TREM2 (chr6), CHCHD10 (chr22), and CCNF (chr16). Long reads establish contiguous, full-length exon-intron connectivity across complex loci where short-read assemblies are fragmented or incomplete.

### Isoform complexity and length distribution

Across the expressed set, we documented a median of 2 isoforms per gene, with 65% resolved as having two or more (Fig. 5a). Isoform lengths spanned a broad range with a median of 2,854 nucleotides (Fig. 5a). This degree of isoform-level resolution is a direct consequence of the read-spanning capability of long reads, which assign a single read to a complete transcript structure irrespective of gene complexity; a distinction that fragment-based short-read assembly cannot reliably achieve^19^.

To illustrate this improved resolution at neurobiologically relevant genes, we examined transcript structures at four loci central to Alzheimer’s disease (AD), Parkinson’s disease (PD), amyotrophic lateral sclerosis (ALS), and frontotemporal dementia (FTD) (Fig. 5c). At GBA1, the most common genetic risk factor for PD and Lewy body dementia ^20,21^, SALRR resolved 4 isoforms with full exon connectivity versus 3 from short-read data, including 3’ variants proximal to the pseudogene GBAP1 that are unresolvable by short reads due to sequence homology. At CCNF, linked to familial ALS and FTD ^22^, SALRR recovered 2 isoforms, including an extended 3’ isoform indistinguishable from noise in short-read data, which detected only 1. At CHCHD10, implicated in ALS, FTD, and PD^23,24^, SALRR resolved 3 isoforms at this complex, overlapping CHCHD10/C22orf15 locus, including one with a substantially extended 3’ region, against 2 truncated isoforms from short reads. At TREM2, a major genetic risk locus for late-onset AD^25,26^, SALRR resolved 5 isoforms exceeding the 3 reference-annotated models, while short-read data recovered only 2. Across all four loci, SALRR recovered transcript structures and isoform ratios partially or completely inaccessible to short-read approaches (Supplementary Table 7), illustrating the disease-relevant transcriptomic complexity that population-scale long-read sequencing can now resolve.

### Long-read sequencing uncovers complex *GBA1* transcript isoforms

Given the technical complexity of the GBA1-GBAP1 locus, we next examined the underlying long-read evidence supporting transcript models resolved by SALRR (Fig. 6). Short-read data cannot reliably resolve transcript architecture here because of extensive sequence identity with the pseudogene GBAP1; long reads instead anchor each isoform to single full-length molecules. We identified two uncataloged GBA1 isoforms, ISOMT_8486 and ISOMT_8487, both showing a novel upstream exon segment and extended 5’ junction boundaries, and both exceeding baseline detection thresholds (median CPM>1) across the 10 brain samples (Fig. 6a; Supplementary Table 8). Split-read alignments confirmed coverage across the novel exon sequences and supporting junction arcs (Fig. 6b, Supplementary Fig. 4), with ISOMT_8486 showing consistently higher expression than ISOMT_8487 across samples (Fig. 6c). These findings indicate that full-length long-read sequencing resolves pseudogene mapping interference at the GBA1 locus and reveals consistently expressed GBA1 transcripts with alternative 5’ structure.

**Fig 6:**
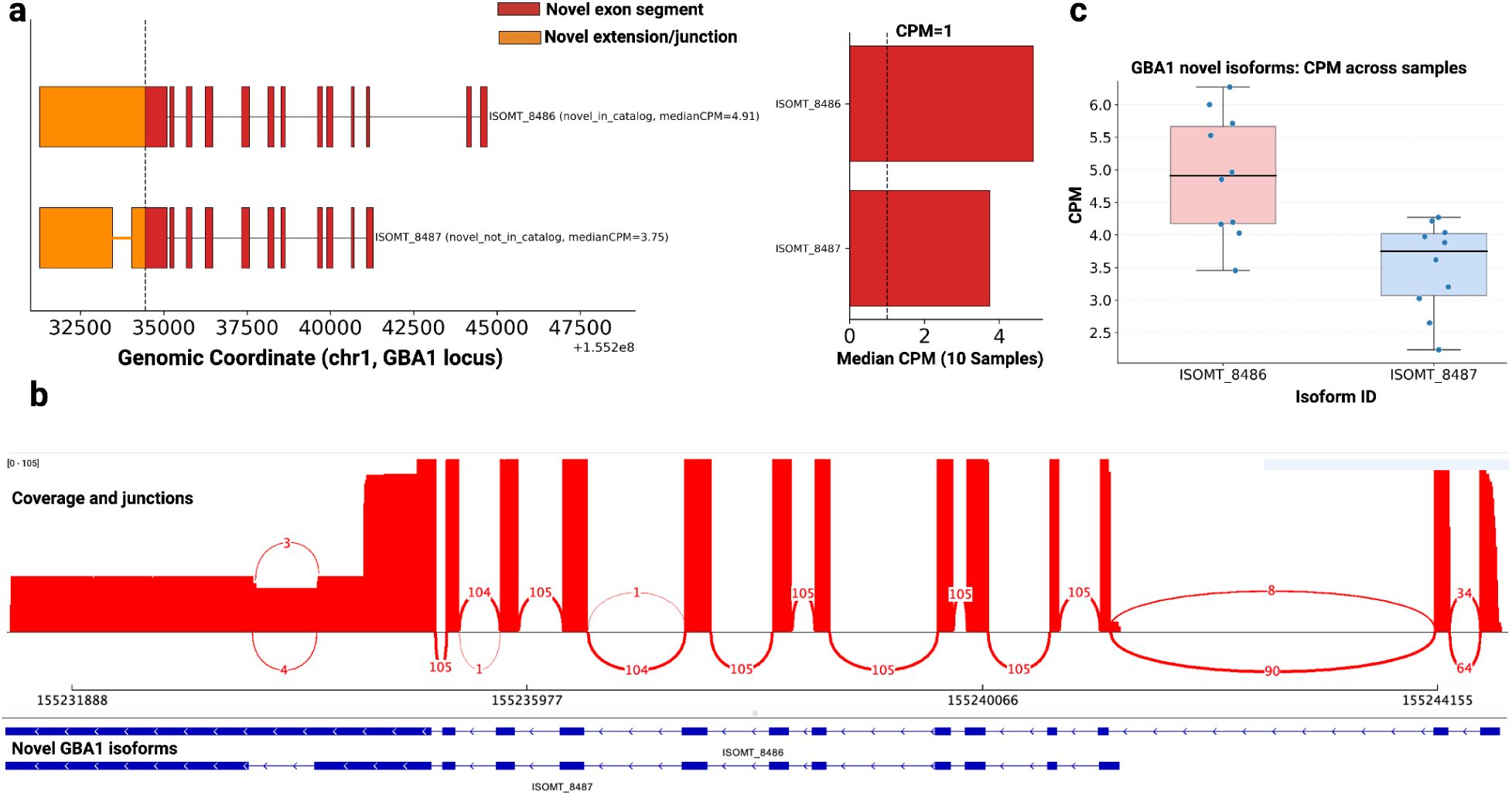
Junction-level evidence of long-read resolved GBA1 transcript isoforms. **a,** Gene models of two novel GBA1 transcripts (ISOMT_8486, ISOMT_8487) at the GBA1 locus on chromosome 1. Red blocks, novel exon segments; orange blocks, novel extensions and junctions relative to the reference annotation. Right: median CPM per isoform across the 10 analysed samples, with the detection threshold at CPM = 1. **b,** Sashimi plot of long-read coverage at the GBA1 locus in one donor (NABEC_SH-97-09). Reads were restricted to those containing the complete internal splice-junction chain of either isoform (±20 bp tolerance), then pooled and deduplicated, giving 105 reads (98 for ISOMT_8486, 7 for ISOMT_8487). Histograms show read depth; arcs show splice junctions labelled with spanning read counts, drawn above or below the axis by read orientation. Lower tracks show the exon–intron structure of both isoforms. **c,** Expression (CPM) of both isoforms across the 10 individual samples.

**Supplementary Fig. 4:**
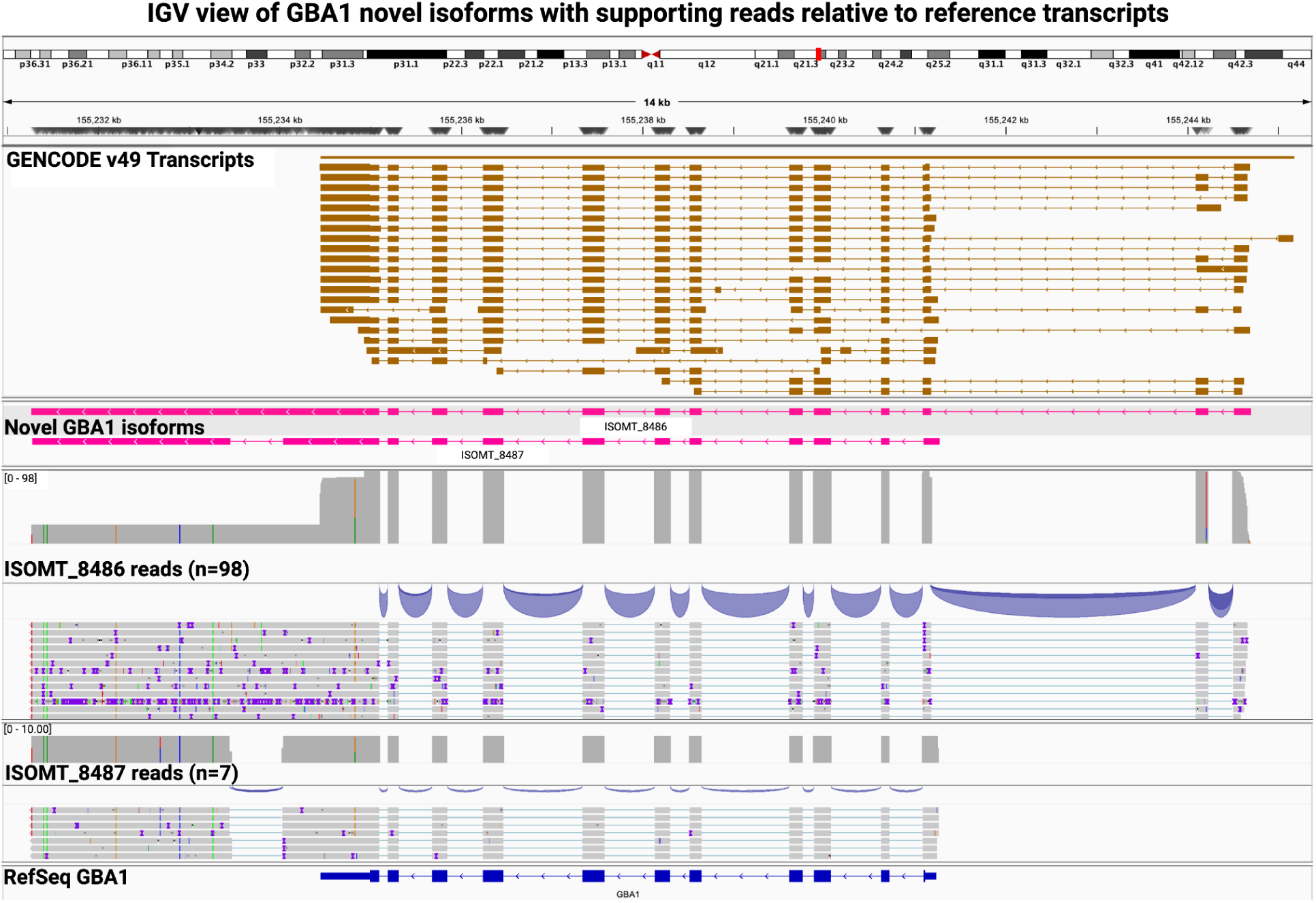
Long-read support for two novel GBA1 isoforms. IGV view of the GBA1 locus (chr1:155,230,948–155,245,358, GRCh38). Tracks from top: GENCODE v49 GBA1 transcripts; the two novel isoforms identified in the 10-brain Isomatch merged annotation, ISOMT_8486 and ISOMT_8487; aligned long reads supporting ISOMT_8486 (n = 98) and ISOMT_8487 (n = 7), each shown with its coverage and splice-junction track; and the RefSeq GBA1 transcript model. Reads are from a single donor (NABEC_SH-97-09, frontal cortex, cDNA) and were retained only if they contained the complete internal splice-junction chain of the corresponding isoform, in order, within a ±20 bp boundary tolerance; end-to-end coverage of the isoform was not required. ISOMT_8487 is defined by a novel intron absent from ISOMT_8486 and from all GENCODE-annotated GBA1 transcripts, and the supporting reads show the corresponding splice gap. Read rows are downsampled by IGV for display; the counts above reflect all qualifying reads.

### Mitochondrial transcript processing

**Supplementary Fig. 5:**
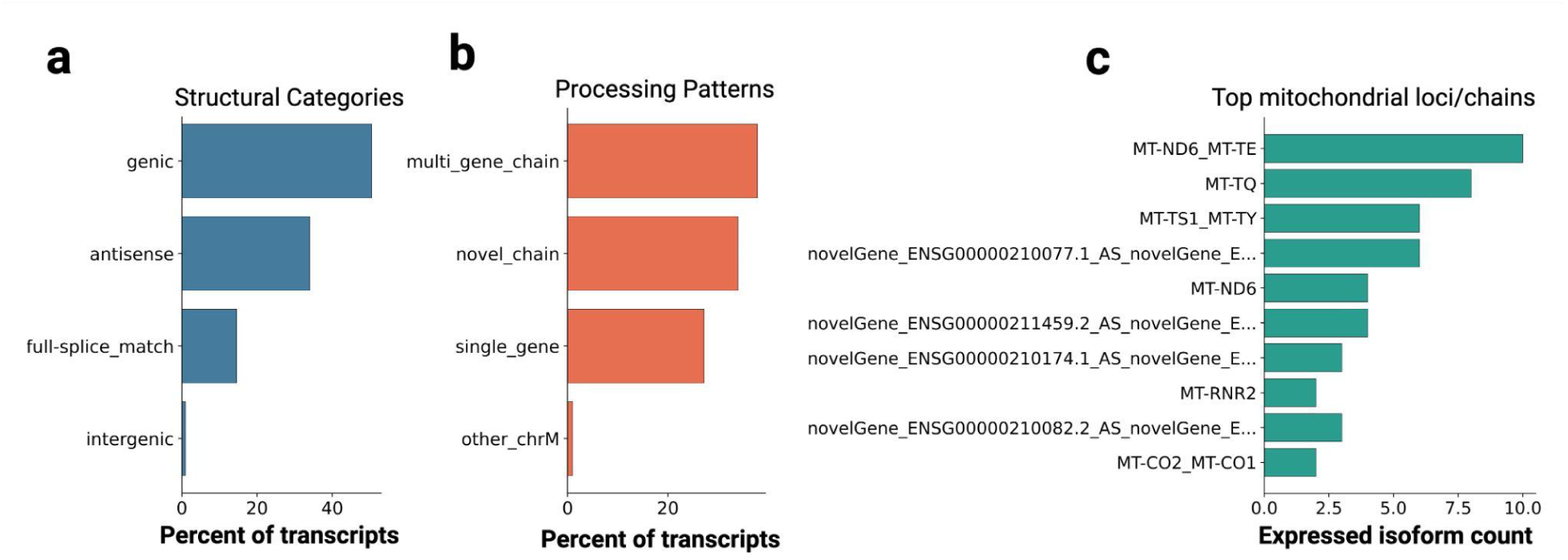
Architectural and processing landscape of mitochondrial transcripts. **a,** Relative percentage distribution of transcript structural categories mapped to the mitochondrial genome. **b,** Breakdown of transcript processing patterns, categorizing transcripts into multi-gene chains, novel unannotated chains, single-gene structures, or other chromosome M regions. **c,** Top 10 most abundant mitochondrial loci or multi-gene chains ranked by their unique expressed isoform count.

Mitochondrial reads were filtered following Aguzzoli Heberle et al.^13^, yielding 103 expressed mitochondrial transcript models across 59 loci (predominantly processing intermediates and readthrough products of polycistronic transcription rather than spliced isoforms). Multi-gene chains spanning adjacent MT genes were most common (∼37%; e.g., MT-ND6_MT-TE), followed by antisense-to-annotated-feature transcripts (∼33%) and single-gene transcripts (∼27%) (Supplementary Fig. 5b); structurally, transcripts were largely genic (∼50%) or antisense (∼33%), with 15% matching annotated splice structures (Supplementary Fig. 5a). The most frequently recovered chain, MT-ND6 and its adjacent tRNA MT-TE (Supplementary Fig. 5c), reflects the light strand’s single protein-coding gene, illustrating the ability of long reads to preserve co-transcribed mitochondrial connectivity inaccessible to short-read assembly. Five mitochondrial isoforms reported by Aguzzoli Heberle et al.¹³ were not detected in our dataset, likely reflecting cohort composition; their study combined AD and control samples (n = 6 each), whereas ours is exclusively healthy controls.

### RNA variant calling in post-mortem human brain

**Supplementary Fig. 6:**
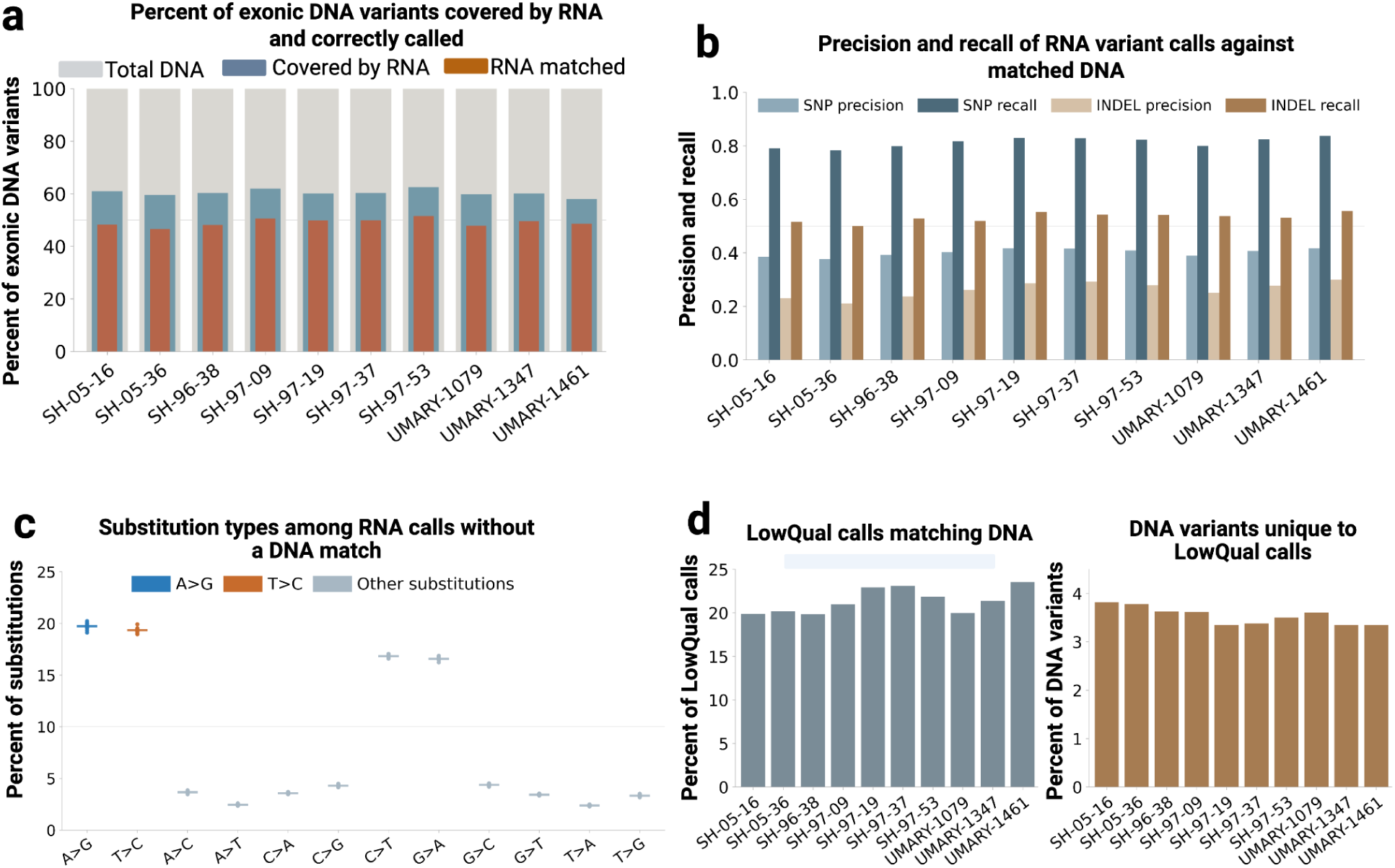
Benchmarking long-read RNA variant calls against matched donor DNA in ten NABEC frontal cortex samples. **a,** Grey, all autosomal SNPs from matched short-read DNA in GENCODE v43 exons outside GIAB difficult regions (100%); blue, the subset covered by at least four RNA reads; orange, the subset Clair3-RNA called correctly. Only 58–64% had sufficient RNA coverage, and 79–82% of those were matched. **b,** hap.py precision and recall for PASS calls over the same region: SNP recall 0.78-0.84, precision 0.38-0.42, with indels lower for both. The DNA truth set was filtered on minor allele frequency, so genuine rare variants in RNA score as false positives and precision is understated. **c,** The twelve substitution types among PASS RNA calls with no matching DNA variant; each dot is one donor, the line the mean. A>G and T>C each reached ∼20% and together 39%, the symmetric pattern expected from ADAR editing on strand-agnostic cDNA. C>T and G>A (∼17%) reflect deamination background; all others fell below 5%. **d,** Left, LowQual (QUAL<8) calls matching a DNA variant (20–24%); right, DNA variants matched only by a LowQual call and not by any PASS call (3.3–3.8%). Note the different y-axis ranges.

Long-read RNA sequencing supports accurate small-variant detection, as recently established by Clair3-RNA across PacBio and ONT direct-RNA platforms^27^, though this has not been examined at population scale in post-mortem human brains. We applied Clair3-RNA to 10 NABEC frontal cortex samples and 3 KOLF2.1J iPSC lines sequenced with ONT R10 cDNA chemistry; brain samples yielded 2.3-fold more variant calls than iPSC (mean 636,580 vs. 278,466), driven largely by REDIportal-tagged editing sites (Supplementary Table 9). Benchmarking against matched short-read genotypes showed that only 58–64% of truth-set SNPs were RNA-callable, of which 79–82% were recovered, indicating that expression depth, not calling accuracy, is the primary constraint on recovery (Supplementary Fig. 6a). Clair3-RNA achieved SNP recall of 0.78–0.84 with precision of 0.38–0.42 (lower for indels; Supplementary Fig. 6b); much of the residual false-positive signal reflects ADAR-mediated RNA editing (A>G/T>C, 39% of RNA-only calls combined) rather than caller error (Supplementary Fig. 6c). Benchmarking RNA variant calling in post-mortem tissue therefore requires a different definition of detectability than reference-line benchmarks assume.

### End-to-end scalability from tissue to isoform-level quantification

Combining the automated wet-lab workflow with SALRR processes 24 samples from frozen tissue to isoform-level quantification tables in 5 days, the time a comparable manual workflow needs for 8 (Fig. 1). This improvement is achieved without sacrificing data quality: mean read depth, mapping rate, isoform detection, and quantification accuracy were equivalent across manually and automatically prepared cohorts processed through SALRR (Fig. 2b). Applied to 10 frontal cortex samples from the NABEC cohort, SALRR produced a complete isoform-resolved transcriptome resource comprising 31,607 unique transcript models and 8,532 novel isoforms, with gene and transcript level quantification matrices ready for downstream statistical analysis.

Previous long-read RNA sequencing studies of the frontal and developing cortex have used manual workflows without integrated analysis pipelines at comparable or larger scale^13,14^. This study does not yet exceed those in sample number, but demonstrates an automated, end-to-end platform designed for population-scale deployment. Because samples are processed concurrently, adding samples does not extend cohort runtime. With automated wet-lab throughput, the platform is ready for immediate application to large postmortem brain cohorts, such as those in the CARD Long-Read Sequencing Initiative (https://card.nih.gov/research-programs/long-read-sequencing), where hundreds to thousands of samples can be processed without modification to the workflow presented here.

## DISCUSSION

Here we present SALRR, an integrated wet-lab and computational platform for cohort-scale, isoform-resolved long-read RNA sequencing from postmortem human brain tissue. It resolves three barriers that have until now made large-scale long-read RNA sequencing in postmortem brain impractical: i) the low throughput and inter-operator variability of manual library preparation; ii) sensitivity of existing protocols to postmortem RNA degradation; and iii) the fragmented, poorly scalable computational landscape for long-read RNA analysis^5,28^. Automation reduces hands-on time by 67% and increases throughput to 24 libraries per operator per day with consistent performance across RINs spanning 8.2 to 10, and SIRV spike-in controls enabling cross-batch calibration^29^. Computationally, SALRR encapsulates basecalling, full-length read identification, reference alignment, dual-tool transcript reconstruction, and isoform-level quantification in a single reproducible workflow, with integrated quality control minimising false-positive transcript calls through multi-stage filtering, stringent mapping thresholds, and SQANTI-like classification^4^. Concurrent processing keeps runtime independent of sample number, making the pipeline immediately deployable on large postmortem brain cohorts.

Our evaluation confirms the workflow captures genuine biological distinctions between iPSC-derived and postmortem brain samples despite technical equivalence between them: library yield, fragment size, Q-scores, and mapping rate were consistent across both sample types despite differing RIN distributions. Against this consistent technical baseline, brain tissue showed a lower proportion of fully annotated transcripts and a correspondingly higher novel isoform and intron-retention burden than iPSC, reflecting the greater cellular and splicing complexity of brain tissue rather than a technical artifact. Polyadenylation profiles reinforced this: mitochondrial poly(A) tails were uniformly short in both sample types, while nuclear poly(A) tails were consistently shorter in iPSC than in postmortem brain, with every iPSC sample falling below the combined median, consistent with the shorter tails of actively cycling cells versus the longer, more stable transcripts of terminally differentiated neurons^30,31^.

The biological findings from the NABEC application illustrate the scale of isoform-level information invisible to short-read transcriptomic studies of neurodegeneration. Across *GBA1, CCNF, CHCHD10,* and *TREM2, SALRR* consistently resolved additional isoforms, extended 3’ ends, and pseudogene-proximal structures that short-read data either missed entirely or could not disambiguate (Fig. 5c). These findings build on a small but growing body of long-read transcriptomic studies in the human brain, including a 10-sample aged frontal cortex resource from Aguzzoli Heberle and colleagues^13^, and a prenatal-to-postnatal cortex atlas from Bamford and colleagues^14^. SALRR extends these contributions in three respects: the automated workflow and demonstrated tolerance to variable RNA integrity make the platform applicable to large, heterogeneous postmortem cohorts rather than the carefully selected samples manual protocols favor; the integrated end-to-end pipeline eliminates the reproducibility and scalability limitations of fragmented multi-tool workflows; and, most consequentially, the generation of matched long-read genomic, epigenomic, and transcriptomic data within the same cohort^32^ creates a framework for multi-omic integration prior studies have not achieved.

Several limitations of the current platform warrant acknowledgement. Long-read cDNA sequencing retains technical biases inherent to ONT chemistry, including homopolymer errors that can affect exon-boundary and UTR assignment, and size-selection biases against very short or long transcripts ^6,7^. SIRV spike-in controls provide cross-batch calibration but do not fully recapitulate endogenous brain RNA complexity ^29^. Postmortem tissue introduces biological variability through agonal state effects and cellular heterogeneity that RNA integrity metrics alone cannot capture ^33,34^ and bulk sequencing cannot resolve cell-type-specific isoform usage, an important caveat given the known cellular heterogeneity of neurodegeneration ^35^. Novel isoform calls are inherently constrained by reference annotation completeness ^4^, and orthogonal validation remains important for confirming novel transcript structures^36^.

The modular architecture of SALRR is designed to accommodate emerging technologies without pipeline redesign; extension to single-cell and spatial long-read RNA sequencing would resolve cell-type-specific isoform usage, a key limitation of the bulk approach presented here^28,35^. SALRR is now being applied to hundreds of AD and related dementia brain samples through the CARD Long-Read Sequencing Initiative, generating isoform-resolved transcriptomic data integrated with matched long-read genomic and epigenomic profiles from the same donors. We anticipate that SALRR will enable transcript-level analyses, including splicing QTL mapping, allele-specific isoform expression, and isoform-switching at disease loci, that have not previously been achievable at this scale in neurodegeneration.

## Methods

### Ethics oversight

The North American Brain Expression Consortium (NABEC) study was approved by the Joint Addiction, Aging, and Mental Health Data Access Committee. Full study details and data access instructions are available on the dbGaP portal under accession number phs001300.v4.p1. In accordance with the U.S. National Institutes of Health (NIH) guidelines, research utilising postmortem human tissue does not constitute human subjects research and did not require additional Institutional Review Board approval.

### Sample collection

Frozen frontal cortex tissue was obtained from *n* = 10 neurologically normal individuals from the NABEC study, which served as a pilot to establish and benchmark the SALRR workflow prior to population scale application. The samples were provided by the Banner Sun Health Research Institute Brain and Body donation program^37^. All donors were of European ancestry with no documented clinical history of neurological, cerebrovascular, or cognitive disorders. Demographic and clinical metadata are provided in Supplementary Table 1. The average age at death was 64 years (range: 19–91 years), with 7 males and 3 females represented.

KOLF2.1J iPSC cells were maintained and frozen as described on protocols.io ^38^. Three frozen pellets of the parental KOLF2.1J line with passage dates in 2022, 2024, and 2025, respectively, were obtained from -80°C storage and processed for RNA extraction.

### RNA extraction and quality control

Detailed protocols for RNA extraction from frontal cortex tissue and cultured cells are publicly available on protocols.io ^39^.

For brain tissue, total RNA was extracted using the RNeasy Lipid Tissue Mini Kit (Qiagen, 74804). Frozen tissue was maintained on dry ice throughout processing. Up to 35 mg of tissue was cut, weighed, and submerged in a custom RNA stabilisation solution for overnight incubation at 4°C. Tissue was homogenized in a TissueLyser III (Qiagen, 9003240) at 25 Hz for 40 seconds using 5 mm stainless steel beads (Qiagen, 69989) and QIAzol lysis reagent. Subsequent steps followed the manufacturer’s protocol, comprising phenol-chloroform extraction, spin column nucleic acid binding, and elution in RNase-free water.

For cultured cells, total RNA was extracted using the RNeasy Plus Mini Kit (Qiagen, 74134). Buffer RLT Plus supplemented with 2-Mercaptoethanol was added to frozen cell pellets (approximately 2 × 10⁶ cells), vortexed, and homogenized through QIAshredder columns (Qiagen, 79656) and gDNA Eliminator columns to remove cellular debris and genomic DNA, respectively, prior to spin column binding and RNA elution according to the manufacturer’s protocol.

RNA concentration was quantified using the Qubit RNA BR Assay Kit (Thermo Fisher Scientific, Q10211) on the Qubit Flex fluorometer (Thermo Fisher Scientific, Q33327), with orthogonal quantification by NanoDrop 8000 spectrophotometry (Thermo Scientific, ND-8000-GL). RNA integrity was assessed using the RNA ScreenTape assay (Agilent Technologies, 5067-5576) on the Agilent 4200 TapeStation system (Agilent Technologies, G2991BA). All samples were stored at -80°C until library preparation.

### cDNA synthesis and library preparation

Complementary DNA (cDNA) libraries were prepared using the cDNA-PCR Sequencing Kit V14 (Oxford Nanopore Technologies, SQK-PCS114). For each sample, 500 ng total RNA was combined with 150 ng Lexogen SIRV Set-4 spike-in controls (Lexogen, 141) to enable cross-batch calibration and assessment of quantification accuracy. Polyadenylated RNA molecules were captured using an adapter incorporating a poly(dT) overhang and reverse-transcribed using Maxima H Minus Reverse Transcriptase (Thermo Fisher Scientific, EP0753). The resulting cDNA was distributed across four parallel PCR reactions to dilute residual reverse transcriptase and minimise amplification bias, then amplified for 22 cycles using Ex Premier DNA Polymerase (Takara Bio, RR370A). cDNA concentration and size distribution were assessed using the Qubit 1X dsDNA HS Assay Kit (Thermo Fisher Scientific, Q33231) and the D5000 ScreenTape assay (Agilent Technologies, 5067-5588), respectively. Libraries were stored at 4°C until sequencing. A minimum of 50 femtomole cDNA was required for adapter attachment and flow cell loading.

### Automated library preparation

To enable population-scale throughput, an automated library preparation protocol was developed on the Hamilton Microlab NGS STAR liquid handling platform using Venus 4 software. The automated workflow executes all steps in a single run, comprising annealing, ligation, digestion, bead cleanup, reverse transcription, PCR amplification, and a second bead cleanup. The primary adaptation from the manual protocol was an increase in input and elution volumes to accommodate well plate geometry, with corresponding adjustments to downstream ligation, reverse transcription, and PCR reaction volumes. Final cleanup elution volumes were similarly increased to ensure complete bead resuspension and minimise sample loss. The automated protocol is publicly available on protocols.io ^39^.

### PromethION sequencing

50 femtomoles of cDNA library were used for rapid adapter attachment. Libraries were loaded onto R10.4.1 flow cells (Oxford Nanopore Technologies, FLO-PRO114M) and sequenced for 72 hours on the PromethION 24 or 48 sequencing system (Oxford Nanopore Technologies, PRO-SEQ024 or PRO-SEQ048). Only flow cells with a starting pore count exceeding 6,500 active pores were used for primary sequencing runs. Sequencing was configured with fast basecalling enabled during acquisition, raw reads output in POD5 format, and a minimum Q score threshold of 8.

### SALRR computational pipeline

All computational processing was performed using SALRR, a modular, Snakemake-based pipeline that processes raw ONT POD5 output through to isoform-level quantification in a single reproducible workflow. The pipeline is publicly available at (https://github.com/NIH-CARD/CARDlongread_ONT_long_read_RNA) and supports deployment on local high-performance computing clusters and cloud environments. A detailed schematic of the full pipeline architecture, including all modules, branching logic, and intermediate file formats, is provided in Supplementary Fig. 1. Key pipeline parameters and software versions are described below and listed in Supplementary Table 3; full configuration files are available in the code repository.

### Throughput and runtime benchmarking

Throughput of the manual and automated library preparation workflows was compared as the number of samples taken from frozen tissue to loaded flow cell within a fixed preparation window (Fig. 1). Computational resource use was profiled on the NIH Biowulf HPC cluster, with per-step CPU and GPU allocations and wall-clock runtimes taken from SLURM job accounting for two technical replicates of a single sample; runtimes exclude scheduler queueing and are reported in Supplementary Table 6.

### Basecalling

POD5 files were basecalled using Dorado v1.2.0 ^40^ with the super-accuracy model (dna_r10.4.1_e8.2_400bps_sup@v5.0.0). Adapter and primer trimming was disabled at the basecalling stage (--no-trim) to preserve full-length read structure for downstream orientation and trimming by Pychopper. Poly(A) tail length estimation was enabled using the --estimate-poly-a flag. Basecalling summary statistics were generated from unmapped BAM output using dorado summary. Additional read-level quality metrics including read counts, length distributions, and output statistics were collected using Cramino v1.1.0 (NanoPack) ^41^.

### Adapter trimming and full-length read identification

Unmapped BAM files were converted to FASTQ format using SAMtools v1.18^42^. Full-length cDNA reads were identified and rescued using Pychopper v2.7.10 ^43^ with the probabilistic hidden Markov model (phmm) classification method. Reads were filtered to a minimum Phred quality score of Q10 and a minimum length of 300 bp. Summary reports and read statistics were generated for downstream visualisation with MultiQC. Post-trimming read quality was assessed using Cramino on the full-length BAM files. An optional contamination screening step using FastQ Screen ^44^ with minimap2 ^45^ alignment against a curated reference database can be enabled or disabled within the pipeline configuration.

### Reference alignment

When SIRV spike-in controls are present (the default configuration), trimmed full-length reads are first aligned to the SIRV genome (SIRV_ERCC_longSIRV_multi-fasta_20210507.fasta) ^29^ using minimap2 v2.28 ^45^ with splice-aware alignment parameters (--splice-flank=no). Reads not mapping to the SIRV reference are extracted and realigned to the human reference genome GRCh38 (GCA_000001405.15; no-alt analysis set). When SIRV spike-ins are not used, SIRV alignment is disabled and reads are aligned directly to the human reference. Alignment quality control was performed at both stages using SAMtools v1.18 ^42^, Cramino ^41^, and mosdepth v0.3.3 ^46^ to capture mapping rates, read length distributions, and coverage depth. Only primary alignments with a minimum mapping quality score (MAPQ) of 40 were retained for downstream transcript reconstruction.

### Poly(A) tail length analysis

Poly(A) tail lengths were estimated by Dorado (--estimate-poly-a) during basecalling and read from the pt:i tag of each aligned read, then partitioned by alignment contig into mitochondrial (chrM) and nuclear sets. Reads with a missing or failed estimate, a zero-length tail, or a value at Dorado’s upper estimation limit (400 nt nuclear, 250 nt mitochondrial) were excluded as uninformative or right-censored. Per-sample distributions were summarised from binned histograms, and figure medians are the median of the thirteen per-sample medians across ten manual brain and three KOLF2.1J iPSC libraries.

### Transcript reconstruction and quantification

Transcript assembly used two complementary tools to maximise sensitivity and specificity. IsoQuant v3.10.0 ^47^ was used for reference-guided isoform detection and quantification with long-read parameters (--data_type nanopore), using its graph-based approach to model complex splicing, assign multi-exon reads to transcript models, and discover novel isoforms alongside GENCODE (.v43) (--genedb, –complete_genedb). In parallel, StringTie v2.2.1 ^48^ was run against the same annotation in long-read discovery mode (-L -G, without -e flag) to reconstruct known and novel transcripts with a minimum coverage threshold of 2.5 (-c 2.5), minimum fraction of 0.05 (-f 0.05), and no end-trimming (-t) to retain high-confidence novel isoforms while filtering assembly noise. Isoforms identified by both tools were merged using Isomatch v0.4.0^49^ against GENCODE v49 annotation^50^, which also classified novel isoforms by structural category, so novel calls are assessed against the current annotation. Quantification outputs were produced at both transcript and gene level to support downstream differential expression and splicing analyses. Per-sample transcript models were structurally classified with SQANTI3 v5.5^36,51^ against GENCODE v43, retaining isoforms passing its default rules-based filter (Supplementary Table 7).

### Cohort merge, requantification, and expression filtering

Per-sample IsoQuant and StringTie models were filtered to remove low-abundance artifacts (CPM > 1) and merged within each sample using Isomatch, as described above. The resulting per-sample consensus models were then merged across the ten brain samples with Isomatch against GENCODE v49 to produce a single unified transcript catalogue for the cohort. Reads from each sample were requantified against this catalogue using IsoQuant, and transcript-level CPM was computed per sample as the transcript read count divided by the total assigned reads, scaled to 10⁶. Transcripts with a median CPM greater than 1 across the ten samples were retained as the high-confidence expressed set, yielding 31,607 transcripts from 10,075 genes. Structural categories for this set were taken from the Isomatch classification against GENCODE v49. The cohort merge, requantification and filtering steps are implemented as pipeline rules within SALRR. Novel transcript models were compared against the high-confidence novel transcript set of Aguzzoli Heberle et al.^13^ using gffcompare v0.12.10 with default parameters, after normalising chromosome names to UCSC style. A model was recorded as overlapping that study if gffcompare assigned it any class code other than *u*, that is any genomic overlap with a transcript in their set.

### Mitochondrial transcript analysis

Mitochondrial transcripts were identified from the Isomatch classification table as isoforms on chrM or with a reference gene name beginning with “MT-”, intersected with the expressed transcript set (median CPM > 1) from the ten manual brain libraries. Each was classed by its match: novel chains (any match involving a novel gene), multi-gene chains (two or more MT-genes), single-gene, or residual chrM. Structural categories were taken from the Isomatch classification, and per-locus isoform counts and summed median CPM were computed by reference gene name. Analyses used pandas, matplotlib and seaborn in Python.

### NABEC bulk short-read RNA sequencing

Bulk short-read RNA-seq for the NABEC frontal cortex cohort was processed as described in Gibbs et al ^52, 32^ under the study accession phs000249.v1.p1. Gene-level expression was quantified with Salmon^53^ against a filtered GENCODE v43 human transcriptome index across 206 frontal cortex samples. For the gene-level correlation analysis, we used existing Salmon TPM estimates, restricted to the 10 donors with matched long-read data. For transcript-level quantification and structure comparisons, we independently assembled short-read transcripts from the STAR-aligned BAMs using StringTie v3.0.1 in discovery mode (−c 2.5, −f 0.05) guided by the GENCODE v43 annotation.

### Long-read and short read quantification benchmarking

Long-read and short-read data from the same ten donors were compared at gene and transcript level, with long-read values taken from the per-sample IsoQuant quantifications. Features were restricted to those with a median TPM greater than 1 in each dataset independently, matched by gene and transcript ID, log₂(TPM + 1) transformed, and assessed by Pearson and Spearman correlation. Gene-level values were correlated across pooled gene–sample pairs (Fig. 3a), while transcript-level values, with short-read TPM taken from StringTie assemblies of the matched data, were averaged across samples before correlation (Fig. 3b).

### Gene body coverage

Gene body coverage profiles were computed with RSeQC v5.0.4^54^ (geneBody_coverage.py) using the full GENCODE v43 annotation converted to BED12 format. Coverage was calculated on genome-aligned, coordinate-sorted BAM files and scaled across 100 equally sized bins spanning each gene from the 5’ to the 3’ end, with per-sample profiles normalised to the maximum bin value. Profiles were generated for three NABEC frontal cortex samples and for the matched short-read alignments from the same donors, and the two sets were overlaid in R.

### Long-read RNA variant calling and benchmarking

RNA variants were called from long-read cDNA alignments using Clair3-RNA^27^ (model ont_r10_dorado_cdna), with REDIportal^55^ tagging of known RNA-editing sites (GRCh38) and the phasing model enabled. Variants passing the caller’s default filters (QUAL ≥ 8, depth ≥ 4, allele fraction ≥ 0.08) were retained as PASS calls. To evaluate calling accuracy, RNA calls were benchmarked against matched short-read NABEC DNA genotypes from the same donors, used as the truth set. Comparisons were restricted to GENCODE v43 exonic regions on autosomes, intersected with each sample’s RNA-callable region (defined as at least four reads of coverage) and excluding the GIAB difficult regions^56^. Concordance was assessed with hap.py^57^ on PASS variants, reporting precision, recall, and F1 for SNVs and indels (Supplementary Fig. 6b), alongside the proportion of truth variants falling inside the RNA-callable region and the proportion of those recovered (Supplementary Fig. 6a). Single-nucleotide false positives, that is PASS calls with no DNA counterpart, were tallied into the twelve base changes as a percentage of each sample’s total (Supplementary Fig. 6c). LowQual records (QUAL < 8) from the unfiltered RNA VCF were matched to the truth set by position and allele over the same intervals (Supplementary Fig. 6d).

### Spike-in recovery analysis

Expected concentrations, lengths and GC content were parsed from the Lexogen SIRV-Set 4 sequence design overview, with poly(A) tails stripped and SIRV isoforms restricted to complete annotations. Observed abundances were transcript-level TPM from StringTie assemblies of the SIRV-aligned reads, matched by GenBank accession for ERCC and by identifiers for SIRV and long SIRV. Because SIRV and long SIRV are supplied equimolar, quantitative accuracy was assessed for ERCC only; the others were scored for detection rate alone. Per sample, ERCC transcripts with non-zero expected and measured values were fitted by linear regression on log_10_ scale, reporting Pearson’s r, R² and Spearman’s ⍴. Accuracy across RNA quality was tested by correlating per-sample ERCC r values against RIN.

### Visualization

Read alignments were visualized in the Integrative Genomics Viewer (IGV) v2.18.2 from genome-aligned, coordinate-sorted BAM files, with the GENCODE annotation loaded as a reference track (Supplementary Fig. 4); the sashimi plot in Fig. 6b was generated in IGV from the same alignments. Transcript structure comparisons in Fig. 5c were drawn with pyGenomeTracks v3.9, plotting the GENCODE v43 reference annotation alongside per-sample long-read IsoQuant and short-read StringTie transcript models, with each locus displayed over the gene span extended by 50 kb on both sides.

### Statistics

Comparisons of library preparation metrics between manual and automated protocols used matched RNA samples and were assessed by two-sided Wilcoxon signed-rank tests. Correlations were quantified using Pearson’s r for continuous relationships and Spearman’s ⍴ where rank agreement was reported. No correction for multiple comparisons was applied. All statistical analyses were performed in Python 3.9.15 using SciPy 1.11.1.

## Code availability

The SALRR pipeline and all associated scripts will be made publicly available upon publication via GitHub (https://github.com/NIH-CARD/CARDlongread_ONT_long_read_RNA/) under an open-source MIT licence, and can be made available to editors and reviewers on request. The repository contains the full Snakemake workflow, conda environment specifications, configuration files, and a quickstart guide to support deployment on local high-performance computing clusters and cloud environments. Additional details about computational tools, software versions, and parameters are provided in the Methods section and in the repository README.

## Data availability

Raw and processed long-read RNA sequencing data generated in this study are available through controlled access in accordance with the data sharing agreements governing the NABEC cohort. Access requires submission of a dbGaP application under accession number phs004923.v1. Demographic and clinical metadata for all samples are provided in Supplementary Table 1. Code to reproduce all figures and analyses presented in this study is available in the GitHub repository listed under Code Availability.

## Acknowledgements

We thank members of the North American Brain Expression Consortium (NABEC) for providing samples derived from brain tissue. We are grateful to the Banner Sun Health Research Institute Brain and Body Donation Program of Sun City, Arizona for the provision of human biological materials. This work was supported in part by the Intramural Research Program (IRP) of the National Cancer Institute (NCI), the National Human Genome Research Institute (NHGRI, ZIAHG200398), National Institute on Aging (NIA, ZIAAG000534, ZIAAG000538), and the Center for Alzheimer’s and Related Dementias (CARD), within the Intramural Research Program of the NIA and the National Institute of Neurological Disorders and Stroke (NINDS). We acknowledge the support of Oxford Nanopore Technologies staff in helping develop the automated cDNA protocol. This work utilized the computational resources of the NIH STRIDES Initiative (https://cloud.nih.gov) through the Other Transaction agreement -Azure: OT2OD032100, Google Cloud Platform: OT2OD027060, Amazon Web Services: OT2OD027852. This work utilized the computational resources of the NIH HPC Biowulf cluster (https://hpc.nih.gov). Some authors’ participation in this project was part of a competitive contract awarded to DataTecnica LLC by the NIH to support open science research. M.A.N. also owns stock in Character Bio Inc. and Neuron23 Inc. This research was supported in part by the IRP of the NIH. The contributions of the NIH authors are considered Works of the United States Government. The findings and conclusions presented in this paper are those of the authors and do not necessarily reflect the views of the NIH or the U.S. Department of Health and Human Services.

## Supplementary tables

**Supplementary Table 1. NABEC demographic and clinical metadata.** One row per sample. Columns: sample ID, age at death, sex, postmortem interval (hours), brain region, RNA integrity number, and ancestry.

**Supplementary Table 2. Per-sample sequencing metrics for all SALRR libraries.** One row per sample. Columns describe library preparation method (manual or automated), sequencing batch, total reads (millions, Q ≥ 8), Median read length (bases), data output, GRCh38 mapping rate (%), median q-score, and number of isoforms and genes detected (before cohort merging and median cpm>1 filtering).

**Supplementary Table 3. SALRR pipeline software versions and parameters.** Complete list of all tools used in the pipeline with version numbers, key parameters applied, and citations. Columns: tool name, version, function in pipeline, key parameters, reference.

**Supplementary Table 4. Novel isoform catalogue.** Full list of novel transcript models identified across the NABEC cohort. Columns: transcript ID, gene name, SQANTI-like structural category, detection counts per sample, source sample IDs, mean expression (TPM), and overlap with Aguzzoli Heberle et al. 2025.

**Supplementary Table 5. Spike-in recovery metrics per sample.** Detection counts against expected totals for SIRV, ERCC, and long SIRV controls from the SIRV-Set 4 mix, alongside ERCC quantitative accuracy (Pearson r, Spearman ⍴, R², and deviation from expected). Columns also record sample ID and library preparation mode.

**Supplementary Table 6. Hardware specifications and computational runtimes.** Step-by-step CPU/GPU resource allocations, sample sequencing metrics (RIN, yield, read count), and execution times across replicate runs on the NIH Biowulf HPC cluster. Columns include sample ID, sequencing yields, hardware resources allocated per step, step-specific runtimes, and total pipeline processing time.

**Supplementary Table 7. GBA1, CCNF, CHCHD10, and TREM2 isoform quantification.** Isoform-level expression values for GBA1, CCNF, CHCHD10, and TREM2 across all 10 NABEC frontal cortex samples. Columns include gene name, transcript ID, SQANTI like structural category, mean expression across the cohort (TPM), standard deviation, proportion of samples in which the transcript was detected, whether the isoform is detectable by matched short-read RNA-seq, and GENCODE release 43 annotation status. Values support the quantitative claims made in Fig. 5c.

**Supplementary Table 8. Novel GBA1 transcript models.** Exon-level coordinates for two novel GBA1 transcripts identified in NABEC frontal cortex by long-read RNA sequencing, provided in GTF format. Models were derived from a cross-sample merge of ten brain samples using Isomatch, aligned to GRCh38 and annotated against GENCODE version 49. Both extend the annotated five-prime end of GBA1.

**Supplementary Table 9. Clair3-RNA variant call filter breakdown.** Per-sample distribution of variant calls by filter status for the ten NABEC frontal cortex samples and three KOLF2.1J iPSC replicates. Columns: sample ID, total variant calls, PASS calls, LowQual calls, calls flagged as putative RNA editing sites, RefCall, other, and the percentage of total calls passing all filters.

## Ethics declarations

## Competing Interests

The authors declare no competing interests.

## Notes

### Competing Interest Statement

The authors have declared no competing interest.

